# Reconstitution of an active human CENP-E motor

**DOI:** 10.1101/2022.01.21.477187

**Authors:** Benjamin Craske, Thibault Legal, Julie P.I. Welburn

## Abstract

CENP-E is a large kinesin motor protein which plays pivotal roles in mitosis by facilitating chromosome capture, alignment and promoting microtubule flux in the spindle. So far, it has not been possible to obtain active human CENP-E to study its molecular properties. *Xenopus* CENP-E motor has been characterised *in vitro* and is used as a model motor, however its protein sequence differs significantly from human CENP-E. Here, we characterise human CENP-E motility *in vitro*. Full-length CENP-E exhibits an increase in run length and longer residency times on microtubules when compared to CENP-E motor truncations, indicating that the C-terminal microtubule binding site enhances the processivity when the full-length motor is active. In contrast to constitutively active human CENP-E truncations, full-length human CENP-E has a reduced microtubule landing rate *in vitro*, suggesting that the non-motor coiled coil regions self-regulate motor activity. Together, we demonstrate that human CENP-E is a processive motor, providing a useful tool to study the mechanistic basis for how human CENP-E drives chromosome congression and spindle organisation during human cell division.

## Introduction

Chromosome alignment and segregation is essential to ensure genomic stability. The mitotic spindle is the physical apparatus that allows the accurate alignment of chromosomes during mitosis. Following the disassembly of the nuclear envelope in prophase, chromosomes are captured by microtubules and aligned in the metaphase plate (Magidson et al., 2015; Pereira et al., 2018; Rodriguez-Rodriguez et al., 2018; Sacristan et al., 2018). However, chromosomes at the spindle poles often cannot biorient through this search and capture mechanism and use a dynein/CENP-E dependent pathway (Bancroft et al., 2014; Barisic et al., 2014). The microtubule motor protein CENP-E is recruited to the fibrous corona of unattached kinetochores, a large macromolecular structure that maximizes the microtubule-binding surface of kinetochores to favour microtubule capture (Cooke et al., 1997; Pereira et al., 2018; Rodriguez-Rodriguez et al., 2018; Sacristan et al., 2018). Upon microtubule capture, CENP-E walks towards microtubule plus ends and promotes the lateral to end-on conversion of kinetochores on microtubules (Shrestha and Draviam, 2013; Sikirzhytski et al., 2018; reviewed in Craske and Welburn, 2020). CENP-E is recruited to kinetochores through a rapid BubR1-dependent and a slower BubR1-independent pathway (Ciossani et al., 2018; Johnson et al., 2004; Legal et al., 2020). Inhibition or depletion of CENP-E in human cells increases the incidence of chromosome misalignments, causes spindle assembly checkpoint activation and results in a prometaphase arrest (Gudimchuk et al., 2013; Tanudji et al., 2004; Wood et al., 2010), highlighting the essential function of the kinetochore-localized motor during chromosome congression. More recently the kinetochore-bound CENP-E population has been implicated in promoting microtubule flux in prometaphase (Steblyanko et al., 2020). CENP-E also localizes to the overlapping microtubules of the spindle midzone and midbody, suggesting roles for CENP-E during the later stages of mitosis (Yen et al., 1991).

Previous work to reconstitute the activity of native CENP-E fractionated from HeLa cells indicated that the full-length protein was inactive (DeLuca et al., 2001). Thus until now, biochemical characterisation studies and *in vitro* reconstitutions of CENP-E activity have used the *Xenopus laevis* CENP-E orthologue. *X. laevis* CENP-E displays processive motility along single microtubules *in vitro* and is required for chromosome alignment in egg extracts (Gudimchuk et al., 2013; Kim et al., 2008; Yardimci et al., 2008). This has provided important insights into how CENP-E functions at a molecular level. However, human and *X. laevis* CENP-E share only 49% sequence similarity. The human model system is often used for cell biology, functional and structural studies of human kinetochores and cell division. Currently it is not clear to what extend the large sequence differences provide properties to human CENP-E distinct from the *Xenopus* CENP-E orthologue to mediate chromosome segregation in humans. In this study, we report that both truncated and full-length human CENP-E motors are active. We find truncated CENP-E is constitutively active and processive *in vitro*, capable of unidirectional movement along microtubules. In contrast, only a fraction of full-length human CENP-E motors are active, yet more processive than truncated CENP-E upon a successful collision with the microtubule. This indicates that the long non-motor region interferes with the motile properties of full-length CENP-E *in vitro*. Overall, the reconstitution of active human CENP-E motors obtained in this study represent a useful ressource for the study of the mechanistic basis for chromosome segregation in humans.

## Results

### Truncated human CENP-E constructs are motile and processive

As human full-length CENP-E has been shown to be inactive (DeLuca et al., 2001), we first tested whether a minimal human CENP-E motor displayed any motility. We designed several N-terminal truncations containing the motor domain. Human CENP-E is predicted to contain over 20 discontinuous coiled-coils within its stalk and C terminus (Fig 1A and 1B). The first putative coiled-coil of human CENP-E is predicted to form between residues 334-401 (Fig 1A) by PairCoil2 (McDonnell et al., 2006). A minimal truncated *Xenopus* CENP-E_1-473_ construct containing the motor domain, a single coiled-coil between residues 335-392 and terminating at Thr-473 with a C-terminal GFP tag, is processive *in vitro* (Kim et al., 2008). We therefore designed a similar construct of human CENP-E, which we refer to as CENP-E_483_, followed by two tandem mNeonGreen fluorophores for recombinant expression and purification from insect cells (Fig 1B and S1B).

**Figure 1.**
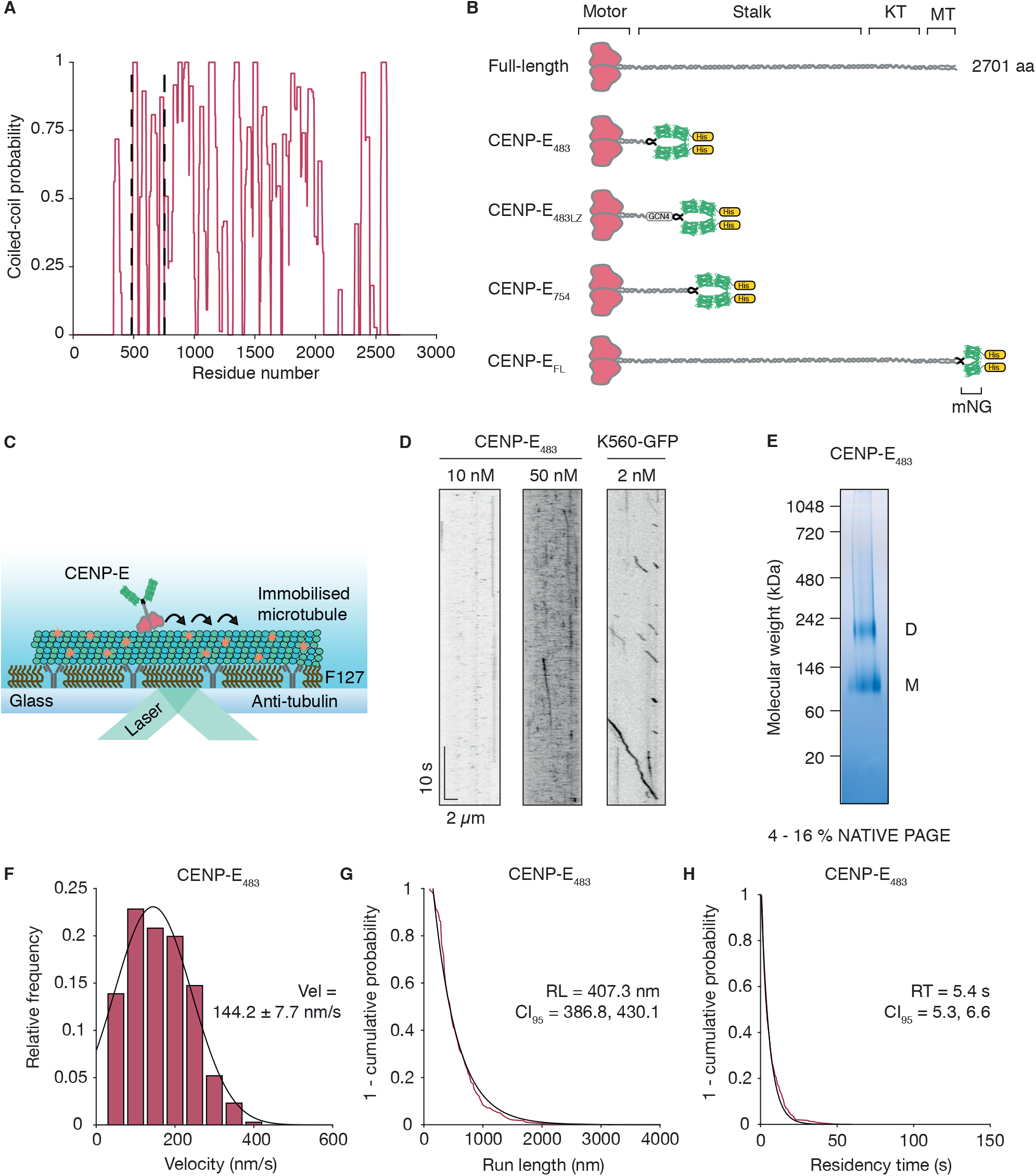
The first predicted coiled-coil of human CENP-E weakly facilitates dimerisation of motor domains. (A) Coiled-coil prediction of full-length CENP-E by Paircoil2. Dashed vertical lines represent truncations. (B) Constructs used in this study. KT = kinetochore binding domain, MT = second microtubule binding site, GCN4 = GCN4 leucine zipper domain, His = hexahistidine tag, mNG = mNeonGreen. (C) Schematic representation of a single molecule motility assay. (D) Kymographs of CENP-E_483_ and K560-GFP at indicated nanomolar concentrations for motility assays. (E) Native PAGE analysis of purified CENP-E_483_ oligomeric status. M = monomer, D = dimer. (F) Histogram representation of velocities for CENP-E_483_ (n=346) at 50 nM fit to a single gaussian distribution (r^2^ = 0.978). (G) 1 - cumulative frequency distribution of run lengths for CENP-E_483_ at 50 nM (n=346) fit to a single exponential decay (r^2^ = 0.982). (H) 1 - cumulative frequency distribution of residency times for CENP-E_483_ at 50 nM (n = 346) fit to a single exponential decay (r^2^ = 0.990).

Next, we tested whether human CENP-E_483_ walks processively on microtubules using *in vitro* reconstitution and single molecule imaging with TIRF (total internal reflection fluorescence) microscopy. Processive landing events of human CENP-E_483_ on immobilized GMPCPP-stabilized microtubules were rare at the low nanomolar concentrations required for single molecule imaging (Fig 1C and 1D), in contrast to constitutively active Kinesin-1 (K560-GFP) (Fig 1D). The majority of CENP-E_483_ microtubule binding events were short-lived interactions, with only a small fraction undergoing continuous unidirectional movement along microtubules (Fig 1D). This suggested to us the motor may not be stable, despite eluting as a single peak by size-exclusion chromatography (Fig S1A). To test whether CENP-E_483_ was a stable dimer, we analyzed the oligomeric status of purified CENP-E_483_ by native PAGE. We detected the presence of two separate protein species migrating at approximately ~100 kDa and ~230 kDa (Fig 1E). Given that the predicted monomeric molecular weight of CENP-E_483_ is 109 kDa, this result indicates that purified CENP-E_483_ exists dynamically as a mixture of monomers and dimers in solution (Fig 1E). Thus the first coiled-coil within the stalk of human CENP-E supports only weak dimerization of the motor, as previously reported for *Xenopus* CENP-E (Yardimci et al., 2008).

We next measured the behaviour of CENP-E_483_ motors *in vitro* at 50 nM, to increase the probability of detecting processive events. We found that human CENP-E_483_ motors exhibited an average velocity of 144.2 ± 7.7 nm/s when moving unidirectionally on the microtubule (Fig 1F). This velocity was approximately 10-fold faster than a previously reported gliding speed for a truncated human CENP-E construct (Sardar et al., 2010). With a run length of 407.3 nm (95% confidence interval, CI_95_ [386.9, 430.1] nm), we found that CENP-E_483_ motors exhibited a relatively long residency time of 5.41 s (95% confidence interval, CI_95_ [5.29, 6.56] s) on the microtubule lattice (Fig 1G and 1H). Kymograph analysis indicated that human CENP-E_483_ often exhibited discontinuous motion and frequently paused during processive runs, with recorded velocities ranging from 16.4 nm/s up to 388.9 nm/s (Fig 1F). Similarly, heterogeneity has also been previously reported in the motility of truncated *Xenopus* CENP-E_1-473_ motors *in vitro* (Gudimchuk et al., 2013; Kim et al., 2008).

### Reconstitution of robust processive motility by human CENP-E through stabilization of a dimeric stalk

To stabilize the CENP-E motor as a dimer, we artificially dimerized truncated human CENP-E construct by fusing a GCN4 leucine zipper domain to the C terminus of CENP-E_483_ and purified CENP-E_483LZ_ (Fig 1B and 2A). This approach has been successful in stabilizing the dimeric state of human KIF1A truncations and reconstituting the superprocessive motility of KIF1A *in vitro* (Soppina et al., 2014). Single molecules of CENP-E_483LZ_ walked along microtubules with an average velocity of 179.9 ± 3.6 nm/s, similar to that measured for CENP-E_483_ (Fig 2B - 2D, Movie 1). However, CENP-E_483LZ_ motors were more processive than the weakly dimeric CENP-E_483_, displaying a run length of 685.2 nm (95% confidence interval, CI_95_ [661.4, 710.7] nm) and a maximum recorded run of 4.4 μm (Fig 2B - E). CENP-E_483LZ_ motors demonstrated residency times of 6.36 s (95% confidence interval, CI_95_ [6.17, 6.56] s) on the microtubule (Fig 2F). Similarly to CENP-E_483_, we found that processive runs of CENP-E_483LZ_ were discontinuous but often included pauses mid-run, leading to longer total residency times on the microtubule before detachment (Fig 2B and 2F).

**Figure 2.**
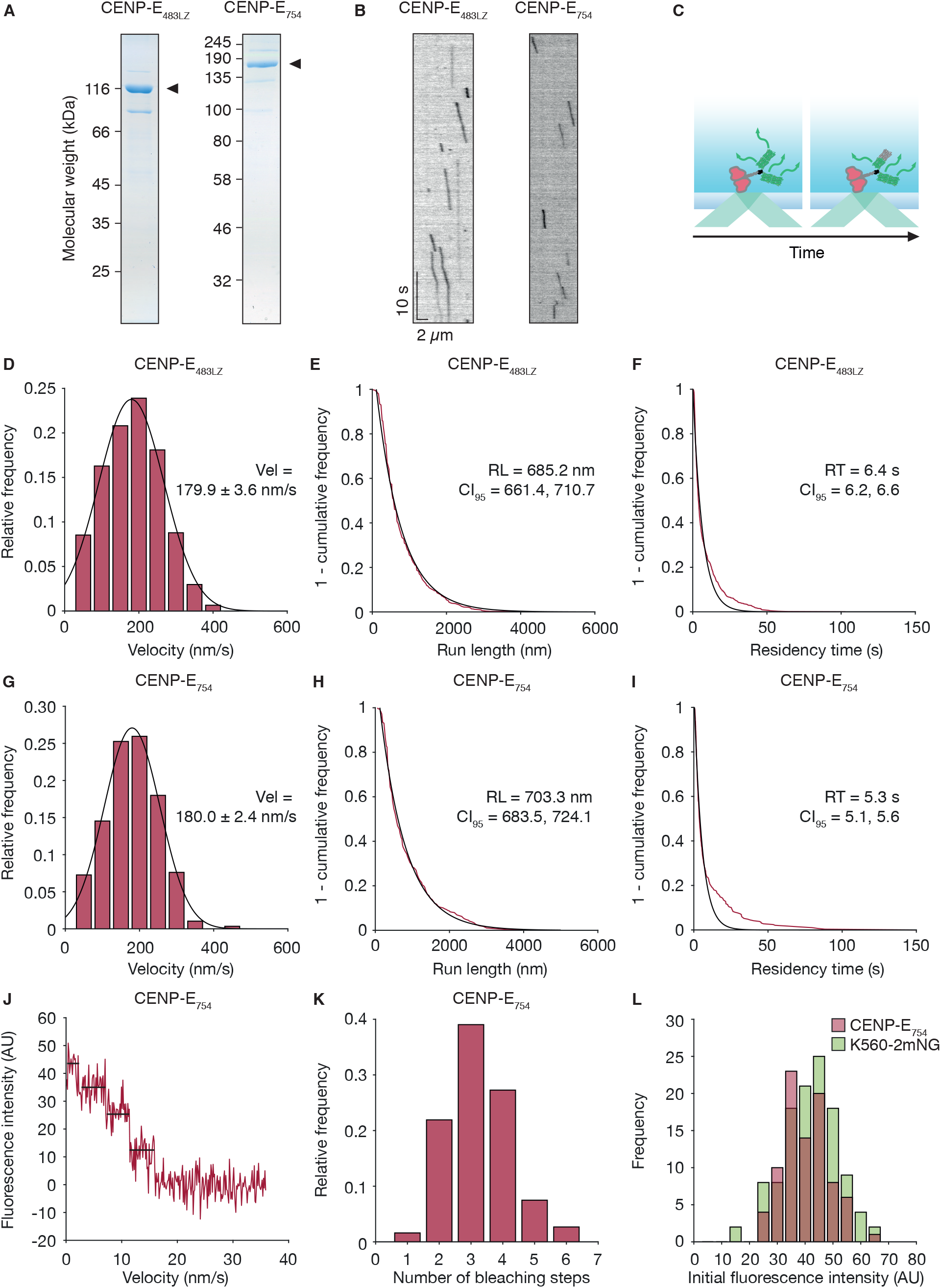
Stable CENP-E dimers are robustly processive in vitro. (A) Coomassie stained gel of purified CENP-E_483LZ_ and CENP-E754 after SDS-PAGE. Arrowheads indicate purified protein. (B) Kymographs of 5 nM CENP-E_483LZ_ and 3.5 nM CENP-E_754_ moving along single microtubules. (C) Schematic representation of photobleaching and intensity analysis assay. (D) Histogram distribution of CENP-E_483LZ_ velocities (n = 774) fit to a single guassian distribution (r^2^ = 0.992). (E) 1 - cumulative frequency of run lengths measured for CENP-E_483LZ_ (n = 774) and fit to a single expontential distribution (r^2^ = 0.986). (F) 1 - cumulative frequency of residency times measured for CENP-E_483LZ_ (n = 774) and fit to a single expontential distribution (r^2^ = 0.969). (G) Histogram distribution of CENP-E_754_ velocities (n = 289) fit to a single guassian distribution (r^2^ = 0.996). (H) 1 - cumulative frequency of run lengths measured for CENP-E_754_ (n = 289) and fit to a single expontential distribution (r^2^ = 0.993). (I) 1 - cumulative frequency of residency times measured for CENP-E_754_ (n = 289) and fit to a single expontential distribution (r^2^ = 0.966). (J) Example 4-step photobleaching trace of CENP-E754. (K) Histogram distribution of CENP-E754 bleaching steps (n=187). (L) Initial fluorescence intensity distribution of CENP-E_754_ (n=88) and K560-2mNG (n=117).

As full-length human CENP-E is a homodimer in solution (DeLuca et al., 2001), we hypothesized that the dimerization was stabilized by the multiple coiled coils within the native stalk region. In line with this, subsequent coiled-coils scored higher in Paircoil2 probabilities than the first coiled-coil 334-401 (Fig 1A). This was confirmed by Alphafold2 which predicts residues 345-399 to be coiled coils, although it does not predict them to dimerize with confidence (Mirdita et al., 2021). We generated a truncated CENP-E_754_ containing five predicted coiled-coils present in the native stalk of CENP-E. We found that CENP-E_754_ was processive, with an average speed of 180.0 ± 2.4 nm/s (Fig 2G and Movie 2), similar to CENP-E_483LZ_ and CENP-E_483_ (Fig 1F and 2D). Thus, the coiled-coils in the stalk of CENP-E stabilize homodimerization of the motor domains and facilitate processivity. Single molecule analysis of CENP-E_754_ on microtubules revealed a run length of 703.3 nm (95% confidence interval, CI_95_ [683.53, 724.11] nm) and a residency time of 5.3 s (95% confidence interval, CI_95_ [5.07, 5.56] s) (Fig 2D – I). Photobleaching assays and intensity analysis indicated that CENP-E_754_ motors typically bleached in 3 or 4 steps and their initial intensities were similar to a purified K560 construct fused to two tandem mNeonGreen tags, referred to as K560-2mNG (Fig 2C and 2J – L). This indicates for the first time that the human CENP-E motor is an active processive motor.

### Full-length human CENP-E is predominantly inactive but becomes processive upon microtubule binding

Previous work reported that the full-length human CENP-E motor purified from human cells is inactive *in vitro* (DeLuca et al., 2001). We expressed and purified full-length human CENP-E-mNeonGreen, referred to as CENP-E_FL_, from insect cells (Fig 3A). The 692 kDa dimeric CENP-E_FL_ eluted as an elongated molecule by size-exclusion chromatography. We next carried out microtubule gliding assays by tethering CENP-E to the coverslip and flowing in free microtubules in solution with ATP, to assess whether CENP-E_FL_ is active. Many microtubules did not glide along the coverslip, despite binding to immobilized full-length motors. Notably, some microtubules bound and pivoted on the surface (Movie 3), as previously published (DeLuca et al., 2001). A subset of microtubules glided along the coverslip surface, with an average velocity of 115.7 nm/s (Fig 3B), in agreement with our single molecule velocities for truncated human CENP-E motors.

**Figure 3.**
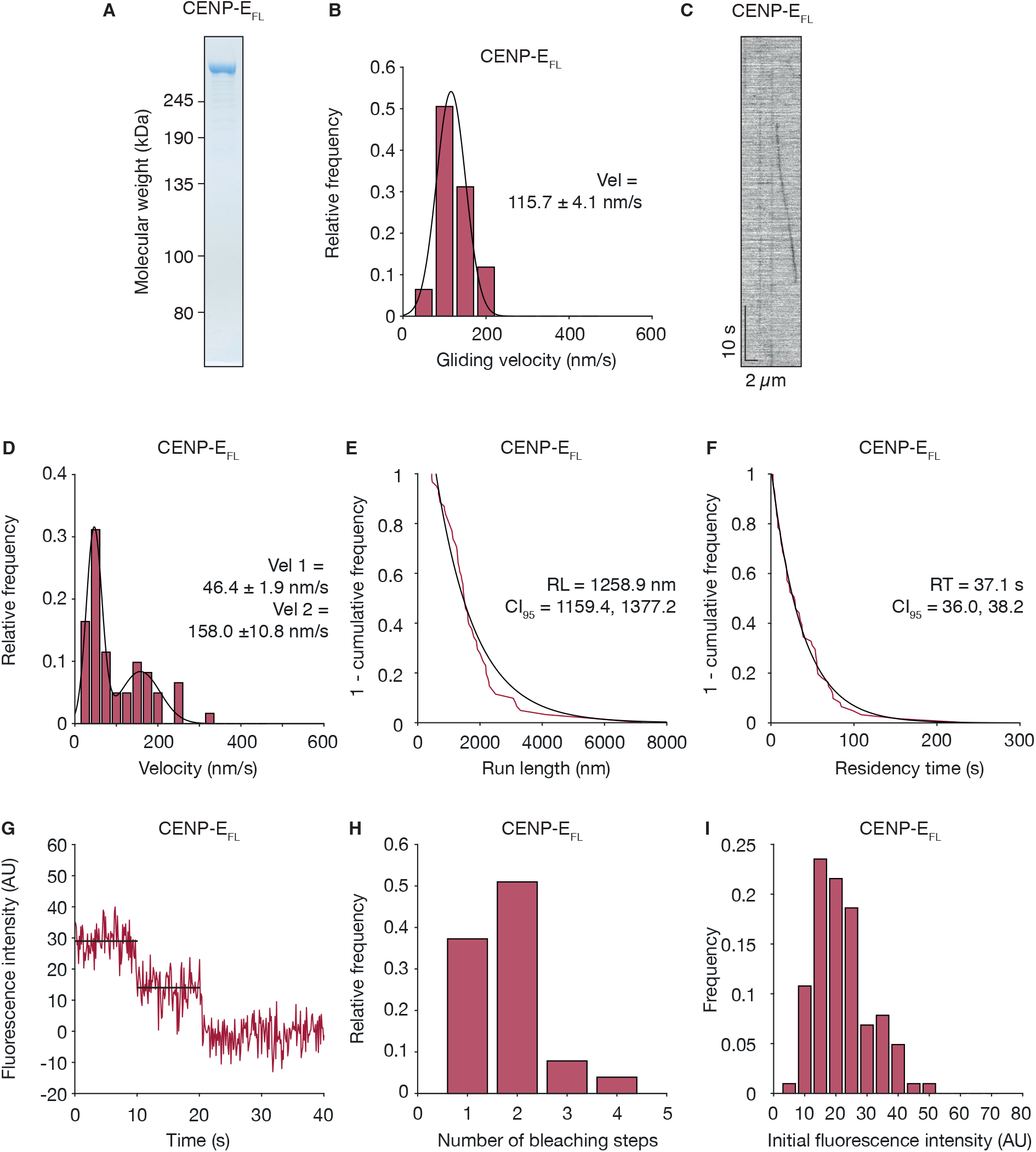
Full-length human CENP-E is a processive motor. (A) Coomassie stained gel of purified CENP-E_FL_ after SDS-PAGE. (B) Histogram distribution for microtubule gliding velocities of CENP-E_FL_ (n = 93). (C) Example of a kymograph showing a single CENP-E_FL_ dimer moving along a microtubule. CENP-E_FL_ was imaged at 12.5 nM. (D) Histogram distribution of CENP-E_FL_ velocities (n = 61) fit to a single guassian distribution (r^2^ = 0.958). (E) 1 - cumulative frequency of run lengths measured for CENP-E_FL_ (n = 61) and fit to a single expontential distribution (r^2^ = 0.951). (F) 1 - cumulative frequency of residency times measured for CENP-E_FL_ (n = 61) and fit to a single expontential distribution (r^2^ = 0.995). (G) Example 2-step photobleaching trace of CENP-E_FL_. (H) Histogram distribution of CENP-E_FL_ bleaching steps (n = 102). (I) Initial fluorescence intensity distribution of CENP-E_FL_ (n = 102).

We next tested whether single CENP-E_FL_ motors displayed any motility on microtubules *in vitro*, using 12.5 nM in our reconstitution assays. We found that CENP-E_FL_ motors predominantly bound to the lattice in a static manner. However, we observed some single molecules moving processively along the microtubule (Fig 3C – I and Movie 4). In contrast to CENP-E_754_, CENP-E_FL_ run length increased 1.8-fold with a run length of 1258.9 nm (95% confidence interval, CI_95_ [1159.42, 1377.22] nm) and a 7-fold increase in residency time to 37.1 s (95% confidence interval, CI_95_ [36.04, 38.15] s) (Fig 3E and 3F). We frequently observed discontinuity in CENP-E_FL_ motion on the microtubule and variation in the recorded velocities (Fig 3D). CENP-E_FL_ displayed a bimodal distribution of velocities and the histogram data were fitted to two overlapping gaussians (Fig 3D). The majority of motile CENP-E_FL_ molecules were slow-moving population of motors, moving at 46.4 ± 1.88 nm/s and would often exhibit paused phases during a single processive run (Fig 3D). Yet, a distinct population of CENP-E_FL_ motors were fast-moving at an average velocity of 157.98 ± 10.77 nm/s, similar to the constitutively active truncated CENP-E constructs characterized above (Fig 2D and 2G). We also found that full-length CENP-E landed on the lattice much less frequently than truncated motors (Fig 4A and 4B). CENP-E_FL_ had a landing rate of 0.147 ± 0.008 events μm^−1^ min^−1^ whereas CENP-E_754_ had a higher landing rate of 0.392 ± 0.008 events μm^−1^ min^−1^ (Fig 4A). Importantly, the processive landing rate of CENP-E_FL_ was 0.009 ± 0.002 events μm^−1^ min^−1^ which was approximately 20-fold lower than the 0.210 ± 0.012 events μm^−1^ min^−1^ observed for truncated CENP-E_754_ motors (Fig 4B). Thus, our *in vitro* reconstitution experiments indicate that a large fraction of purified full-length CENP-E molecules are not active in our assay conditions. However, purified full-length human CENP-E molecules that are active, are highly processive upon a successful collision with the microtubule (Fig 4C).

**Figure 4.**
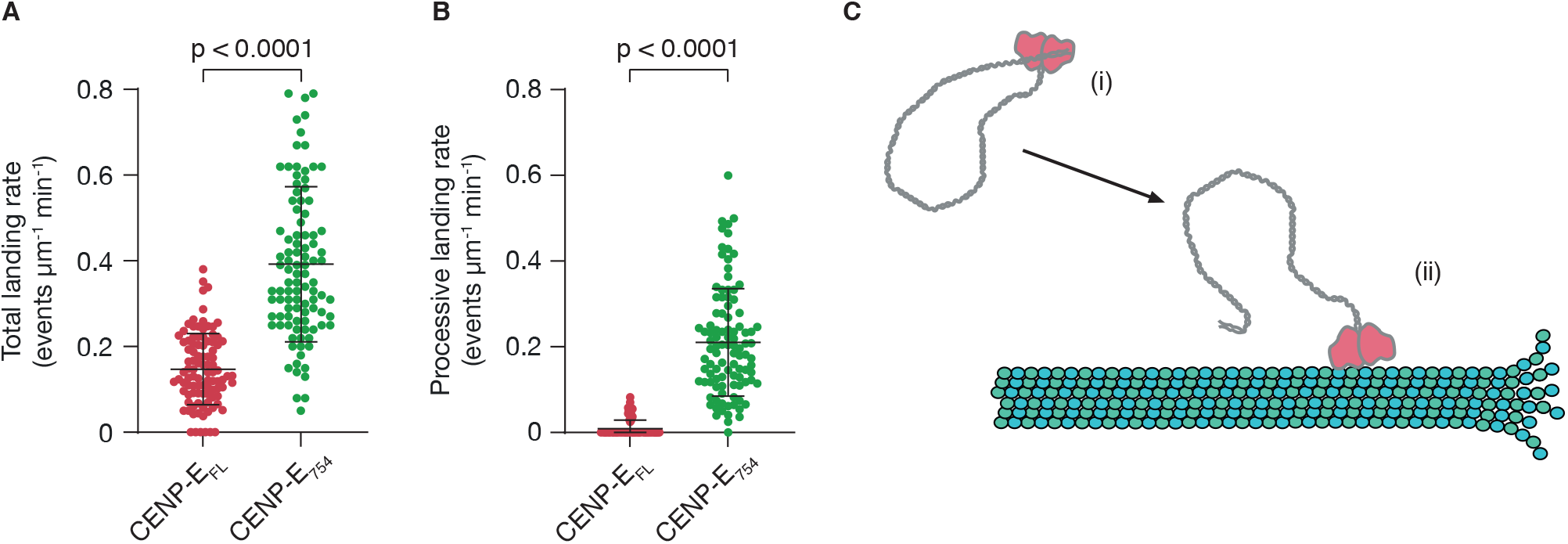
The non-motor regions of human CENP-E regulate processive motility. (A) Quantification of total landing rates for CENP-E_FL_(n = 98, n = number of microtubules) and CENP-E_754_ (n = 100, n = number of microtubules). Welch’s t-test, p < 0.0001. (B) Quantification of processive landing rates for CENP-E_FL_ (n = 98, n = number of microtubules) and CENP-E_754_ (n = 100, n = number of microtubules). Welch’s t-test, p < 0.0001. (C) Model of full-length CENP-E regulation. (i) In the absence of activating cargoes or post-translational modifications, CENP-E does not productively interact with microtubules and processive motility is blocked by autoinhibition. (ii) Once activated, CENP-E is capable of long-lasting processive motility required for transport of chromosomes in prometaphase, sorting of spindle microtubules and stabilisation of end-on kinetochore microtubule attachments.

## Discussion

Taken together, we show that the human CENP-E motor is an active and processive plus-end directed motor. The majority of full-length CENP-E motors move at a slow average velocity of 46.4 ± 1.88 nm/s, with a fraction moving at a comparable velocity to constitutively active truncated motors (Fig 2D, 2G and 3D). Similar behaviour has been previously reported for kinesin-1, whereby motile full-length KIF5B molecules exhibit discontinuity in their processive motion and display a much slower velocity than the KIF5B tail-truncated mutant (Friedman and Vale, 1999). Processive full-length CENP-E motors exhibited higher run lengths and residency times than truncated CENP-E motors. Here we show this is due to the increased stabilization of the dimer through the extensive coiled coils when compared to truncated CENP-E_483_. The increase in processivity may also be due to the presence of a non-motor microtubule binding site at the far C terminus of full-length CENP-E, as CENP-E_754_ lacks this region (Liao et al., 1994; Musinipally et al., 2013; Welburn, 2013). Many kinesins have a second non-specific microtubule binding site which increase their residency time and processivity (Mayr et al., 2011; McHugh et al., 2018; Zhernov et al., 2020).

Our observation that full-length CENP-E activity is relatively variable, may explain why previous attempts to reconstitute microtubule gliding activity of HeLa extract purified CENP-E were unsuccessful (DeLuca et al., 2001). The coiled coils of full-length CENP-E may increase the conformational entropy of the motor *in vitro* and interfere with microtubule binding and processivity. *In vitro* reconstitutions with full-length *Xenopus* CENP-E indicate that the fraction of active motor is increased when coupled to a bead *in vitro*, indicating a potential mechanism where engagement of the C terminus interacting with a cargo (i.e. the bead in that study) promotes CENP-E motor activity (Gudimchuk et al., 2013). We propose interacting partners at the outer corona or the kinetochores could reorganize the coiled coils regions, stabilize an active conformation of CENP-E and co-ordinate its processive transport activity similarly to activation of other kinesin motors (Cho et al., 2009; Coy et al., 1999; Henrichs et al., 2020; Hooikaas et al., 2019; McKenney et al., 2014). Several proteins have been described to interact with CENP-E at kinetochores including BubR1, CLASP1/2, PP1 and CENP-F (Chan et al., 1998; Ciossani et al., 2018; Kim et al., 2010; Kurasawa et al., 2004; Maffini et al., 2009). Super-resolution imaging of kinetochores indicates CENP-E has a compact conformation at the outer corona and kinetochores, close to Ndc80, CENP-F and Spindly (Varma et al., 2013; Wan et al., 2009). *In vitro*, full-length *Xenopus* CENP-E under load stalls at an average force of 4.6 pN but surprisingly maintains a short length of 45 nm when transporting beads under the application of a sidewards force (Gudimchuk et al., 2018). Thus activated CENP-E may maintain a compact conformation during transport of heavy-load cargoes, which includes pulling of chromosomes towards the equator (Kapoor et al., 2006) and potentially sliding cross-linked microtubules of the spindle (Risteski et al., 2021; Steblyanko et al., 2020).

Overall human CENP-E appears to be a less efficient motor than the *Xenopus* orthologue of CENP-E. Here we show that truncated human CENP-E has an average velocity of 179.9 ± 2.4 nm/s and a typical run length of 703.3 nm. Truncated *Xenopus* CENP-E_473_ was first reported as a slow motor with an average speed of 8 nm/s (Kim et al., 2008). However, subsequent reconstitutions with truncated *Xenopus* CENP-E473 and full-length *Xenopus* CENP-E demonstrated 50-fold higher velocities of approximately 300 nm/s and 400 nm/s respectively, and average run lengths between 1.5 – 2.5 μm (Barisic et al., 2015; Gudimchuk et al., 2013). The presence of the C-terminal microtubule binding site in full-length *Xenopus* CENP-E was not reported to enhance CENP-E processivity, in contrast to what we observe for human CENP-E (Fig 2D, 2G and 3D) (Gudimchuk et al., 2013). These discrepancies could be attributed to species divergence. For example, human and *Xenopus* CENP-E proteins share only 49% sequence similarity across their entire length. *Xenopus laevis* CENP-E is 253 residues longer than human CENP-E, with a large insertion C-terminal to the kinetochore-targeting domain. It is also likely we are missing regulatory partners that would stabilize the coil-coiled and kinetochore-binding region of CENP-E to optimize motor activity. Recent studies have highlighted previously unappreciated localization patterns of human CENP-E at overlapping microtubule bundles and to the detachable fibrous corona in human cells (Pereira et al., 2018; Sacristan et al., 2018; Steblyanko et al., 2020). Whether *Xenopus* CENP-E is also recruited to these subcellular regions, or whether this is a human-specific CENP-E function, is not currently known. Given that CENP-E interacts with multiple partners in distinct locations, it will be important to define how the regulatory partners regulate CENP-E structure and function, and how they can affect the load-bearing capacities of CENP-E to fulfil its mitotic functions.

## Experimental procedures

### Protein expression and purification

The sequences for the CENP-E-mNeonGreen gene were made synthetically for this study and are deposited on addgene. mNeonGreen gene was synthesized by Genewiz. Three synthetic DNA fragments of human CENP-E, codon optimised for insect cell expression, were ordered from Gen9. Each DNA fragment contained 100 bp of overlapping fragments and were amplified by PCR and purified. DNA was transformed into competent BY4741 *Saccharomyces cerevisiae* as described in (Gietz and Schiestl, 2007) using an equimolar ratio of each 3 fragments and pRS415 vector, previously linearised with SmaI. Briefly, PEG, Lithium Acetate and herring sperm DNA were incubated with the DNA to be assembled and added to 50 μL of competent cells. After a 30-minute incubation at 30°C, DMSO was added and the cells were heat shocked at 42°C. The cells were then spun down, re-suspended in 400 μL of 5 mM CaCl_2_ and plated on Synthetic Defined medium without Leucine. Genes encoding full-length *Homo sapiens* CENP-E were amplified by PCR and inserted into a pFastBac1 vector backbone, with a 3C PreScission Protease cleavage site, mNeonGreen fusion protein and a hexahistidine tag located at the C terminus. Truncated *Homo sapiens* CENP-E constructs were generated by PCR amplification of the codon optimised CENP-E sequence as a template. PCR products were digested ligated into a pFastBac1 vector containing 2x tandem mNeonGreen fusion proteins and a hexahistidine tag at the C terminus. K560-2mNG was generated by PCR amplifying the *Homo sapiens* KIF5B sequence (amino acids 1-560) and inserting into a pET3aTR vector (Tan, 2001) containing 2x tandem mNeonGreen fusion proteins and a hexahistidine tag at the C terminus.

Recombinant human CENP-E proteins were expressed using the baculovirus system in Sf9 cells. Cells were harvested 48-62 hours after infection and stored at −70 °C until use. Harvested cells were resuspended in CENP-E lysis buffer (50 mM HEPES pH 7, 300 mM KCl, 40 mM imidazole, 1 mM MgCl_2_, 1 mM EGTA, 0.1 mM ATP, 5 mM betamercaptoethanol) supplemented with 1 mM PMSF, 5 μg/mL DNase and 1x cOmplete protease inhibitor tablet per 50 mL. Cells were lysed in a dounce homogeniser with 30-40 strokes. The lysate was cleared by centrifugation at 40,000 rpm in a Type 45 Ti rotor for 60 minutes at 4 °C and applied onto a pre-equilibrated HisTrap HP column (GE Healthcare) in CENP-E lysis buffer at 4 °C. HisTrap columns were washed with 40 column volumes of CENP-E lysis buffer. Proteins were eluted with 250 mM imidazole. Elution fractions were concentrated, centrifuged at 13,300 rpm for 15 mins at 4°C and then loaded onto a Superose 6 Increase 10/300 column (GE Healthcare) pre-equilibrated with CENP-E gel filtration buffer (50 mM HEPES pH 7, 300 mM KCl, 1 mM MgCl_2_, 1 mM EGTA, 0.1 mM ATP, 1 mM DTT). Fresh CENP-E proteins were used for all *in vitro* motility assays due to deterioration in activity after freezing.

*Homo sapiens* K560-GFP was purified using a previously described protocol (Case et al., 1997) omitting the final microtubule bind and release step, snap frozen and stored at −70 °C. K560-2mNG was transformed in *E. coli* BL21 CodonPlus (DE3) RIL (Agilent Technologies). Transformed BL21 cells were grown to OD_600_ = 0.6 then cooled to 20 °C before induction with 0.5 mM IPTG for 3-4 hours at 20 °C. Frozen pellets were resuspended in K560 lysis buffer (50 mM Tris pH 7.5, 300 mM KCl, 40 mM imidazole, 1 mM MgCl_2_, 1 mM EGTA, 0.1 mM ATP, 5 mM beta-mercaptoethanol) supplemented with 1 mM PMSF and 1x cOmplete protease inhibitor tablet per 50 mL, and sonicated. The lysate was cleared by centrifugation at 58,000 *g* for 50 minutes at 6 °C in a JA25:50 rotor. The supernatant was incubated with Ni-NTA beads (Thermo) for 1.5 hours at 4 C. Beads were washed with 40 column volumes of K560 lysis buffer and proteins were eluted with 250 mM imidazole. Elution fractions were concentrated and loaded onto a Superose 6 Increase 10/300 column (GE Healthcare) preequilibrated with K560 gel filtration buffer (50 mM Tris pH 7.5, 300 mM KCl, 1 mM MgCl_2_, 1 mM EGTA, 0.1 mM ATP, 1 mM DTT). Fractions containing K560-2mNG were snap frozen with 10% glycerol and stored at −70 °C.

### TIRF microscopy

Microscopy was performed on a Zeiss Axio Observer Z1 TIRF microscope using a 100 × NA 1.46 objective equipped with a Photometrics Evolved Delta EMCCD camera and controlled by Zen Blue 2.3 software. For single molecule experiments a 1.6x tube lens was used. The environmental chamber was incubated at 30 °C for all experiments. Coverslips used for motility assays were silanised as in (McHugh et al., 2018). Flow chambers were prepared by attaching a silanised coverslip to a microscopy slide with double-sided sticky tape. 4 sample flow chambers were constructed per microscopy slide, each with a volume of 7-8 μL. Rhodamine microtubules were captured using a 561 nm laser with 15 % intensity, 75 ms exposure. Images of mNeonGreen and GFP tagged motors were captured using a 488 nm laser with 50% intensity, 100 ms exposure and a frame rate of 0.12 frames per second.

For all *in vitro* motility experiments, 0.2 mg/mL GMPCPP (Jena Biosciences) microtubule seeds containing 7% rhodamine-tubulin (Cytoskeleton Inc., TL590M-B,) were polymerised in BRB80 (80 mM PIPES pH 6.9, 1 mM EGTA, 1 mM MgCl_2_) for 1 hour at 37 °C, followed by centrifugation at 13,300 rpm for 10 minutes and then resuspended in BRB80. For gliding assays, anti–His tag antibodies (Raybiotech, 168-10481) at a 1:10 dilution in BRB80 were first introduced to the chamber. Next, 40 μL of 1 % Pluronic F-127 (Sigma Aldrich) in BRB80 was washed through the chamber and incubated for 5 minutes. 200 nM of purified kinesin motors were then added to the chamber in BRB80 supplemented with 2 mM ATP. Chambers were then washed with 1 mg/mL casein (Sigma Aldrich) before a 1:25 dilution of GMP-CPP microtubules was added in final assay mix (80 mM PIPES pH 6.9, 5 mM MgCl_2_, 2 mM ATP, 1 mM DTT and an oxygen scavenger mix: 0.2 mg/ml glucose oxidase, 0.035 mg/ml catalase, 4.5 mg/ml glucose, and 0.1 % beta-mercaptoethanol).

For single molecule motility assays, anti–β-tubulin antibodies (Sigma-Aldrich, T718) at a 1:10 dilution in BRB80 were first introduced to the chamber. Next, 40 μL of 1 % Pluronic F-127 in BRB80 was washed through the chamber and incubated for 5 minutes. GMPCPP microtubules were diluted 1:50 in BRB80 and then added to the chamber for 5 minutes. Chambers were then washed with 1 mg/mL casein (Sigma Aldrich). Freshly purified motors were then added in final assay mix at concentrations indicated in the figure legends and chambers sealed with nail varnish. For photobleaching and intensity analysis, 0.5 nM of fluorescently tagged motor was added to silanised coverslips and allowed to non-specifically adsorb to the surface. After 3 minutes, BRB80 supplemented with oxygen scavenger mix was flown through the chamber to wash away non-adsorbed motors. The sample chamber was imaged using the same conditions as described for single molecule assays.

### Image processing and analysis

Kymographs were manually generated in ImageJ (Schneider et al., 2012). Gliding velocities, single molecule velocities, run lengths and residency times were measured from these kymographs. Data are collected from at least two independent experiments using motors from separate protein purifications. Histograms were generated from raw velocity data and fit to a gaussian distribution in MATLAB (Mathworks). For run lengths and residency times determination, cumulative frequency distributions were generated using the ecdf function in MATLAB and fit to a single exponential distribution. Run length and residency times were represented by the decay constant. Landing rates were determined at a concentration of 3.5 nM for each dimeric motor. Welch’s t-tests were carried out in Graphpad Prism (GraphPad Software). Aggregates as determined from their initial intensity were excluded from analysis. Where necessary, images were also corrected for stage drift using the ImageJ Manual Drift Correction plug-in.

A custom ImageJ macro was developed for analysis of photobleaching steps (https://github.com/bcraske/ImageJ). Briefly, individual fluorescent spots adsorbed to coverslips were manually selected using the multi-point tool in ImageJ. Next, a 4 × 4 pixel square was assigned to the ROI and the average intensity was measured over time using the plot Z axis profile function. Background fluorescence was subtracted by assigning a 10 × 10 square centred around the ROI, excluding the 4 × 4 pixel area, using the plot Z axis profile function. Discrete photobleaching steps were manually counted from the plotted results. Initial intensity values were calculated as the average fluorescence intensity from a single adsorbed motor over the first 5 frames of imaging and the data was plotted as a histogram in MATLAB.

## Acknowledgements

JW is supported by a Wellcome Senior Reseach Fellowship (207430). JW is also a EMBO Young Investigator. B.C. is supported by the Biotechnology and Biological Sciences Research Council (BBSRC) [grant number BB/M010996/1]. The Wellcome Centre for Cell Biology is supported by core funding from the Wellcome Trust (203149).

## Author contributions

JW designed the project. BC performed experiments with CENP-E. TL designed and assembled the CENP-E gene into pFastBac plasmid from gblocks optimized for insect cell expression. BC and JW performed data analysis and interpretation. BC and JW wrote the manuscript.

## Figure legends

**Figure S1. CENP-E_483_ elutes as a single peak during size exclusion chromatography.** (A) Elution profile of CENP-E_483_ from size exclusion chromatography. CENP-E_483_ eluted as a single peak marked by an asterisk (*). (B) Coomassie stained gel showing purified CENP-E_483_ after SDS-PAGE.

**Movie 1. Single molecule motility of CENP-E_483LZ_.** *In vitro* reconstitution of CENP-E_483LZ_ motility. Time lapse imaging of CENP-E_483LZ_ was overlayed onto a single reference snapshot of GMPCPP-stabilised rhodamine microtubules taken from the same field of view, as a reference for microtubule localisation. Scale bar = 2 μm.

**Movie 2. Single molecule motility of CENP-E_754_.** *In vitro* reconstitution of CENP-E_754_ motility. Time lapse imaging of CENP-E754 was overlayed onto a single reference snapshot of GMPCPP-stabilised rhodamine microtubules taken from the same field of view, as a reference for microtubule localisation. Scale bar = 2 μm.

**Movie 3. Microtubule binding, pivoting and gliding by CENP-E_FL_.** Behaviour of GMPCPP-stabilised rhodamine microtubules on a coverslip immobilised with CENP-E_FL_. Scale = 4 μm.

**Movie 4. Single molecule motility of CENP-E_FL_.** *In vitro* reconstitution of CENP-E_FL_ motility. Time lapse imaging of CENP-E_FL_ was overlayed onto a single reference snapshot of GMPCPP-stabilised rhodamine microtubules taken from the same field of view, as a reference for microtubule localisation. Scale bar = 2 μm.

